# The *Candida albicans* Cdk8-dependent phosphoproteome reveals repression of hyphal growth through a Flo8-dependent pathway

**DOI:** 10.1101/2021.06.10.447844

**Authors:** Jeffrey M. Hollomon, Zhongle Liu, Scott F. Rusin, Nicole P. Jenkins, Allia K. Smith, Katja Koeppen, Arminja N. Kettenbach, Lawrence C. Myers, Deborah A. Hogan

**Affiliations:** Department of Microbiology and Immunology, Geisel School of Medicine at Dartmouth, Hanover, NH 03755; Department of Biochemistry and Cell Biology, Geisel School of Medicine at Dartmouth, Hanover, NH 03755

## Abstract

Ssn3, also known as Cdk8, is a member of the four protein Cdk8 submodule within the multi-subunit Mediator complex involved in the co-regulation of transcription. In *Candida albicans*, the loss of Ssn3 kinase activity affects multiple phenotypes including cellular morphology, metabolism, nutrient acquisition, immune cell interactions, and drug resistance. In these studies, we generated a strain in which Ssn3 was replaced with a functional variant of Ssn3 that can be rapidly and selectively inhibited by the ATP analog 3-MB-PP1. Consistent with *ssn3* null mutant and kinase dead phenotypes, inhibition of Ssn3 kinase activity promoted hypha formation. Furthermore, the increased expression of hypha-specific genes was the strongest transcriptional signal upon inhibition of Ssn3 in transcriptomics analyses. Rapid inactivation of Ssn3 was used for phosphoproteomic studies performed to identify Ssn3 kinase substrates associated with filamentation potential. Both previously validated and novel Ssn3 targets were identified. Protein phosphorylation sites that were reduced specifically upon Ssn3 inhibition included two sites in Flo8 which is a transcription factor known to positively regulate *C. albicans* morphology. Mutation of the two Flo8 phosphosites (threonine 589 and serine 620) was sufficient to increase Flo8-HA levels and Flo8 dependent activity, suggesting that Ssn3 kinase activity negatively regulates Flo8. Previous work has also shown that loss of Ssn3 activity leads to increased alkalinization of medium with amino acids. Here, we show that *FLO8* and *STP2*, a transcription factor involved in amino acid utilization, are required for *ssn3*Δ/Δ phenotype, but that loss of the Ssn3 phosphosites identified in Flo8 was not sufficient to phenocopy the *ssn3*Δ/Δ mutant. These data highlight the spectrum of processes affected by the modulation of Ssn3 activity and underscore the importance of considering Ssn3 function in the control of transcription factor activities.

## Introduction

One of the important roles of the Mediator transcriptional co-regulatory complex is to link the activity of promoter-bound transcriptional factors to the basal transcription machinery. The Cdk8 sub-module of Mediator plays important roles in modulating the activity of Mediator itself as well as the activity of transcription factors, among other proteins. In *Candida albicans*, like in other eukaryotes, the Cdk8 module consists of four subunits: the catalytic subunit Ssn3 (Cdk8), Ssn8 (CycC), Med12 (Srb8), and Med13 (Srb9). Ssn3, the catalytic component of the Cdk8 module, is a cyclin-dependent like kinase and its activity depends on the cyclin-like protein Ssn8. Across eukaryotic species, the Cdk8 kinase has been shown to be particularly important for regulation during metabolism and morphpology (1, 2). In mammals, for example, glycolysis, lipogenesis and immune responses are influenced by Cdk8 phosphorylation of specific regulators (3–5). In *S. cerevisiae*, Ssn3(Cdk8) has been well-studied for its roles in metabolism (6, 7) and its negative regulation of pseudohyphal growth through Ste12 (8) and Phd1 (9, 10). The Cdk8 module is of particular interest for its role in transitions between growth conditions and during development as cells need to rapidly make coordinated changes to the abundances and activities of cellular regulators.

In *C. albicans*, null mutations in either *SSN3* or *SSN8* result in interrelated, pleiotropic phenotypes. *SSN3* null mutants had increased induction of drug resistance elements (11) and increased resistance to the effects of bacterially-produced metabolic inhibitors (12). The *ssn3*Δ/Δ leads to a hyperwrinkled colony morphology, increased respiratory metabolism and amino acid utilization (12), and fraction of Ras1 in its active GTP-bound state (13). Mutation of *SSN3* was found to unmask an alternative filamentation pathway in macrophages (14). Recent work by Lu and colleagues (15) found that changes in Ssn3 activity in response to CO_2_, mediated by the Ptc2 phosphatase, led to decreased Ssn3 phosphorylation and decreased inhibition of Ume6, a positive regulator of hyphal growth.

Here, we utilize analog-sensitive variants of *C. albicans* Ssn3, as has been performed in *S. cerevisiae* and in human cells, (3, 16), to study the immediate effects of inhibition of Cdk8 kinase activity on *C. albicans*. Under conditions that do not promote hyphal growth in wild-type strains, Ssn3 inhibition led to the formation of hyphae. We used these conditions to elucidate the *C. albicans* Cdk8 regulon as it related to the control of morphology using phosphoproteomic and transcriptomic analysis of cells shortly after Ssn3 inhibition. Flo8, a transcription factor that positively regulates hyphal growth (17, 18), was identified in the phosphoproteomics analysis as a candidate for Ssn3 regulation, and transcriptomics data showed alterations in hypha-specific genes known to be regulated by Flo8. Deletion of Ssn3-phosphosites in Flo8 was sufficient to affect protein levels and morphology. Additional assays suggest that Flo8 plays major roles in Ssn3-regulated control of metabolism. The data in this manuscript indicate the spectrum of proteins that are altered, directly or indirectly, by Ssn3 kinase activity as cells respond to changing environments. While these studies focus on Flo8, our data show that other transcription factors, including Efg1 and Eed1 are also likely Ssn3 targets and thus the tools and insights presented here may aid in the study of diverse proteins involved in morphology, virulence and drug resistance.

## RESULTS

### 3-MB-PP1 inhibits analog-sensitive Ssn3

To investigate the direct targets of the *C. albicans* Ssn3 kinase, we sought to develop a strain in which Ssn3 kinase activity could be rapidly inhibited using approaches that have been successfully applied in *S. cerevisiae* (16). Based on the alignment of the *C. albicans* and S*. cerevisiae* Ssn3 orthologs, we predicted a phenylalanine to glycine substitution at position 257 would yield a variant that retained the functions of the wild-type kinase, while still being able to be specifically inhibited by the ATP analog 3-MB-PP1 (19). The Ssn3^F257G^ was constructed and is henceforth referred to as the <u>a</u>nalog-<u>s</u>ensitive variant, Ssn3^AS^.

We first assessed the inhibition of Ssn3^AS^ by 3-MB-PP1 with an *in vitro* kinase assay using purified Mediator complex containing the Cdk8 module. Ssn3^WT^ or Ssn3^AS^ were expressed in a background that contained a His-FLAG-tagged derivative of Ssn8 for purification. The activity of Ssn3^WT^ or Ssn3^AS^ was assessed using ^32^P *in vitro* kinase assays (20) in which phosphorylation of recombinantly produced C-terminal domain (CTD) of RNA Pol II, an Ssn3 substrate, was monitored. The Ssn3^WT^ and Ssn3^AS^ kinases had equal CTD phosphorylation activity in the absence of inhibitor (**Fig. 1**). The Ssn3^AS^ kinase activity was inhibited by 3-MB-PP1 in a dose-responsive fashion, while addition of this compound had no effect on the Ssn3^WT^ kinase activity (**Fig. 1**). A concentration of 2.7 μM 3-MB-PP1 inhibited ~85% of the Ssn3^AS^ activity and 24 μM virtually eliminated the activity.

**Figure 1.**
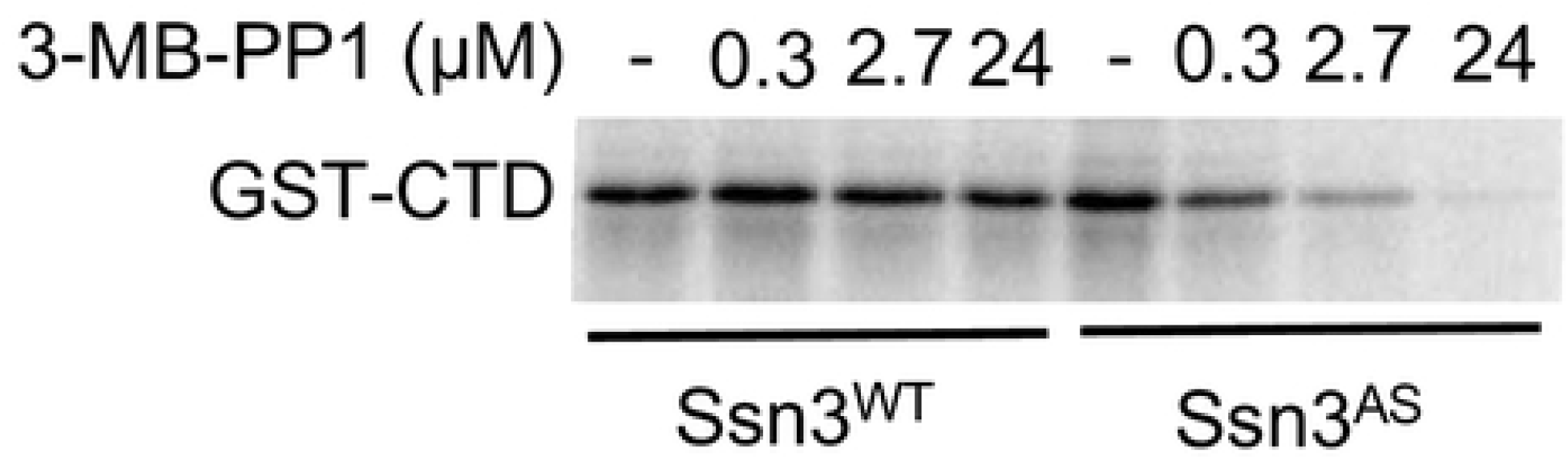
3-MB-PP1 inhibits the activity of analog-sensitive Ssn3^AS^, but not Ssn3^WT^ *in vitro*. *In vitro* kinase reactions contained purified Mediator from a strain with Ssn3^WT^ or Ssn3^AS^, ^32^P-ATP, purified GST-tagged RNA Pol II C-terminal domain (CTD) and the indicated concentrations of 3-MB-PP1 inhibitor. Reactions were analyzed by SDS-PAGE and visualized by phosphorimaging.

In order to determine if Ssn3 kinase activity could be inhibited *in vivo,* we engineered a derivative of *C. albicans* SC5314 in which both alleles of *SSN3* had been replaced by *ssn3^AS^*. We also constructed a strain with two copies of the *ssn3-D325A* allele, which encodes a non-functional, or kinase dead, Ssn3 variant (21) due to mutation of a key aspartate in the catalytic site. This kinase-dead variant is referred to here as *ssn3^KD^*. We have previously shown that, under certain growth conditions, *ssn3*Δ/Δ mutants form hyperwrinkled colonies compared to the SC5314 wild type (WT) (12), a phenotype associated with increased hyphal growth. Thus, we asked if the Ssn3^AS^-expressing strain had a hyperfilamentous phenotype specifically in the presence of 3-MB-PP1 and not in its absence. To quantify hypha formation differences in WT, *ssn3^AS^*, and *ssn3^KD^* strains, we identified conditions which revealed differences in the propensity for hypha formation among strains. Thus, we grew cells in YNB containing amino acids and N-acetyl-glucosamine (GlcNAc), both of which are inducers of hyphal growth (YNBAG), but incubated cultures at 30°C, a temperature lower than that generally used to induce robust hyphal growth. All cultures were amended with either 3-MB-PP1 or the vehicle DMSO. The morphology of the WT was almost entirely yeast and pseudohyphae, and the relative fractions of these morphologies were unaffected by 5 μM 3-MB-PP1 (**Fig. 2A**). While the *ssn3^AS^* cells were similar to the WT in vehicle control cultures, the addition of 3-MB-PP1 to *ssn3^AS^* cultures caused a significant increase in the number of hyphae (**Fig. 2**). The increase in the fraction of cells in the hyphal morphology in 3-MB-PP1 treated *ssn3^AS^* was concomitant with a significant decrease in the fraction of cells present as yeast (**Fig. 2A**). The *ssn3^KD^* strain formed significantly more hyphae than the WT and *ssn3^AS^* strains in control conditions, and the fraction of cells as hyphae was unaffected by the addition of 3-MB-PP1 (**Fig. 2A–B**). Together, these data indicated that 5 μM 3-MB-PP1 inhibits Ssn3^AS^ *in vivo* and that the ATP analog has no discernable effects on the morphology of WT or *ssn3^KD^* strains.

**Figure 2.**
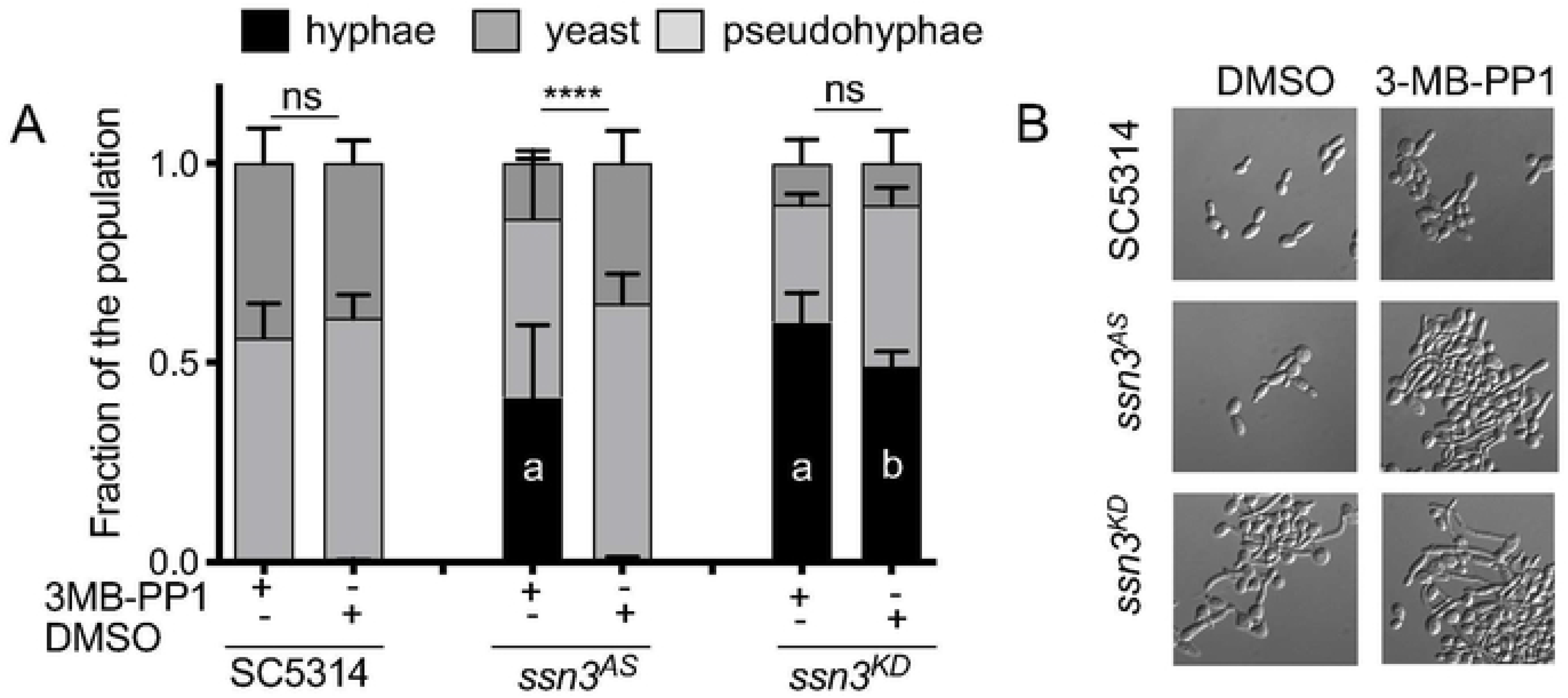
3-MB-PP1 stimulates hyphal growth in a strain bearing analog-sensitive alleles of *SSN3.* **A.** Morphology of wild type (WT) SC5314, *ssn3^AS^* and *ssn3^KD^* strains was assessed after growth in either 5 μM 3-MB-PP1 or vehicle (DMSO) for 3h at 30°C. Quantification of yeast, pseudohyphae and hyphae in cultures by microscopic analysis of blinded samples. ANOVA with multiple comparisons for the hyphal cell populations shown. ****, p<0.001, ns, not significant; a, p<0.001 for comparison to WT with 3MB-PP1; b, p<0.001 comparison to WT with DMSO. **B.** Representative images of cell populations from cultures analyzed in panel A.

### Transcriptomics analyses reveal inhibition of Cdk8 by 3-MB-PP1 leads to the induction of hypha-specific genes

To further explore the effects of Ssn3 inhibition on the regulation of hypha formation (12) and on transcription more broadly, we examined the transcriptomes of *ssn3^AS^* and WT strains grown with 3-MB-PP1 or DMSO vehicle. The WT strain was included in this experiment to assess off-target effects of 3-MB-PP1. Cells from stationary phase cultures were inoculated into the same medium used for morphology assessment, YNBAG, with either 5 μM 3-MB-PP1 or an equivalent volume of DMSO. Three replicate cultures for each strain-treatment combination were incubated for 60 min at 30°C prior to RNA harvest and sequencing as described in the methods. In the *ssn3^AS^* strain, we found that 249 genes were significantly up regulated upon treatment with 3-MB-PP1 by 2-to 49-fold (p<0.05 FDR); transcripts for 33 genes were significantly lower by more than 2-fold with treatment (**Table S2**). Fewer genes were affected by 3-MB-PP1 in the WT with 157 and 4 genes significantly increased and decreased, respectively; the magnitudes of the changes were also smaller (2-to 6-fold) (**Table S2**). The greater number of transcripts at higher levels upon inhibition of Ssn3 is consistent with the Cdk8 module being a predominantly negative regulator of transcription factors (3) or the transcriptional re-initiation or “pausing” of RNA polymerase II (reviewed in (22)). We found seventy-seven transcripts that exhibited a significant fold increase (>2-fold) in both the comparison of the *ssn3^AS^* strain treated with 3-MB-PP1 to its treatment with DMSO, and the comparison of 3-MB-PP1-treated *ssn3^AS^* strain to 3-MB-PP1-treated WT (**Fig. 3** with gene expression shown in log2-transformed counts per million). Consistent with phenotypic analysis of the effects of Ssn3 inhibition on morphology, transcripts encoding *ECE1* and *HWP1* were two of the most highly induced by Ssn3 inhibition in the AS strain (46- and 22-fold higher, respectively). Other transcripts differentially-increased upon Ssn3 inhibition included genes associated with hyphal morphology, such as *ALS1*, *ALS3*, *IHD1*, *RBT1*, *HYR1*, and *HGC1* (14, 23). Consistent with the previous finding that Ssn3 represses induction of Mrr1 controlled genes, *MDR1* was significantly induced with Ssn3 inhibition (11).

**Figure 3.**
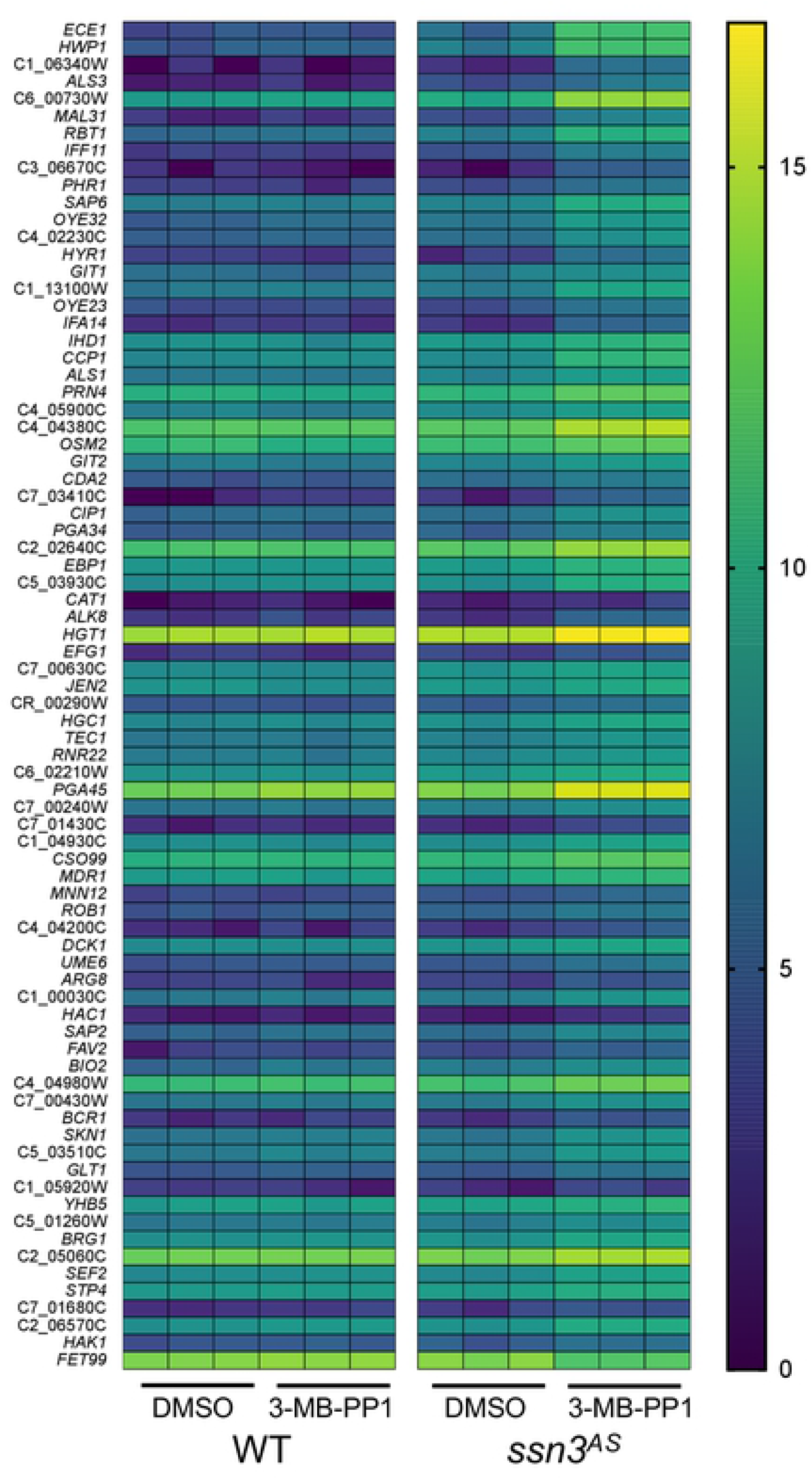
Heat map for genes induced upon inhibition of Ssn3^AS^ by 3-MB-PP1. Seventy-seven genes were 2-fold higher in both the comparison of Ssn3^AS^ with 3-MB-PP1 compared to WT with 3-MB-PP1 and of Ssn3^AS^ with 3-MB-PP1 compared to with the DMSO control. The heat map shows Log2-transformed counts per million for expressed transcripts.

### Specific inhibition of Ssn3 affects the phosphoproteome during the induction of hyphal growth

To identify direct phosphorylation targets of Ssn3 that led to changes in phenotype and the transcriptome, we used mass spectrometry to quantitatively analyze the phosphoproteomes of the *ssn3^AS^* strain compared to the WT after 3-MB-PP1 treatment. We grew *ssn3^AS^* and WT cells to stationary phase, then incubated culture aliquots with 5 μM 3-MB-PP1 for five minutes to allow for drug entry into the cell. We then added concentrated, fresh, pre-warmed YNBAG medium to reproduce the conditions that induce hyphal growth in strains with low Ssn3 activity but not in the WT. Cultures were incubated for an additional 15 min at 30°C with shaking followed by rapid harvest in order to minimize secondary effects of Ssn3 inhibition. Cells were lysed under liquid nitrogen by grinding. We conducted three replicate experiments across three different days, each from an independent overnight culture to increase the robustness of our experimental design.

To reveal the direct targets of Ssn3, we focused on phosphosites that became less abundant in the presence of 3-MB-PP1 in the *ssn3^AS^* strain, but not the WT (**Table 1** and **Table S3** for complete dataset). We found that 977 phosphopeptides mapping to 552 proteins were depleted in *ssn3^AS^* compared to WT, setting a 2-fold cutoff in addition to the P<0.05 test for significance (**Table 1**). Within this group of depleted phosphosites, 82.7% were serines, 15.8% were threonines, and 1.5% were tyrosines. These proportions are very similar to those we observed with a quantification of the whole *C. albicans* phosphoproteome (20), with a slight increase in the number of phosphoserines and a concomitant decrease in the number of phosphothreonines. Although promiscuous, Cdks have been described as proline-directed kinases, and consistent with this, 293 phosphopeptides contained prolines in the position adjacent to C-terminal side of the phosphoresidue, referred to as the Proline Motif (**Table 1**) (24). Fewer phosphosites were found within common RXS/T motifs, in which arginine is in the −2 position relative to the phosphoserine or phosphothreonine, (25). We observed that 264 phosphopeptides (202 proteins) were significantly enriched upon Ssn3 inhibition. This increase in phosphorylation likely represents indirect or secondary effects of Ssn3 inhibition.

**Table 1.** Summary of phosphopeptides and phosphoproteins detected by motif. Analysis limited to those peptides to which P-values of <0.05 were assigned in the comparison of *ssn3^AS^* to SC5134 wild type (WT). Down indicates phosphopeptides lower in *ssn3^AS^* treated with 3-MB-PP1 compared to WT treated with 3-MB-PP1; up indicates phosphopeptides that were more abundant in the *ssn3^AS^* strain treated with 3-MB-PP1 compared to WT treated with 3-MB-PP1. The total refers to the number of peptides or proteins that are significantly different regardless of fold-difference. “#” indicates the detected phosphorylation on a serine or threonine, S/T, in either a −1 position relative to a proline, S/T#P motif.

|  | Down ><br>2-fold | Up ><br>2-fold | Total<br>Down | Total<br>Up |
| --- | --- | --- | --- | --- |
| Total phosphopeptides | 977 | 264 | 2967 | 2522 |
| Total phosphoproteins | 552 | 202 | 1165 | 1057 |
| S/T#P motif phosphopeptides | 293 | 94 | 943 | 867 |
| S/T#P motif phosphoproteins | 218 | 77 | 505 | 493 |

We found 218 proteins that contained phosphosites covered by 2 or more depleted phosphopeptides (a consideration taken to increase stringency) in the *ssn3^AS^* strain compared to WT, and of these 218 proteins, 40 were annotated as having known or predicted nuclear localization in UniProt **(Fig. 4)**(26). Among these, we found that Ssn3AS inhibition led to depletion of phosphopeptides from Med4, with phosphorylated S21 and S33, which is notable as these have previously been identified as target sites for the *C. albicans* Ssn3 kinase (20). We also observed depletion of specific phosphosites (T589 and S620) in Flo8, a regulator of filamentous growth in both *C. albicans* and *S. cerevisiae* (17, 18, 27), and both T589 and S620 were found within the aforementioned proline motif (**Table 1**). Our transcriptomic data showed that *FLO8* levels were unchanged by either Cdk8 inhibition or 3-MB-PP1 addition (**Table S2**). While we focus on Flo8 in the studies presented here, it is worth noting that other hyphal growth associated transcription factors, such as Efg1 were found among the proteins with phosphosites that were at lower abundance with Ssn3 inhibition as discussed below.

**Figure 4.**
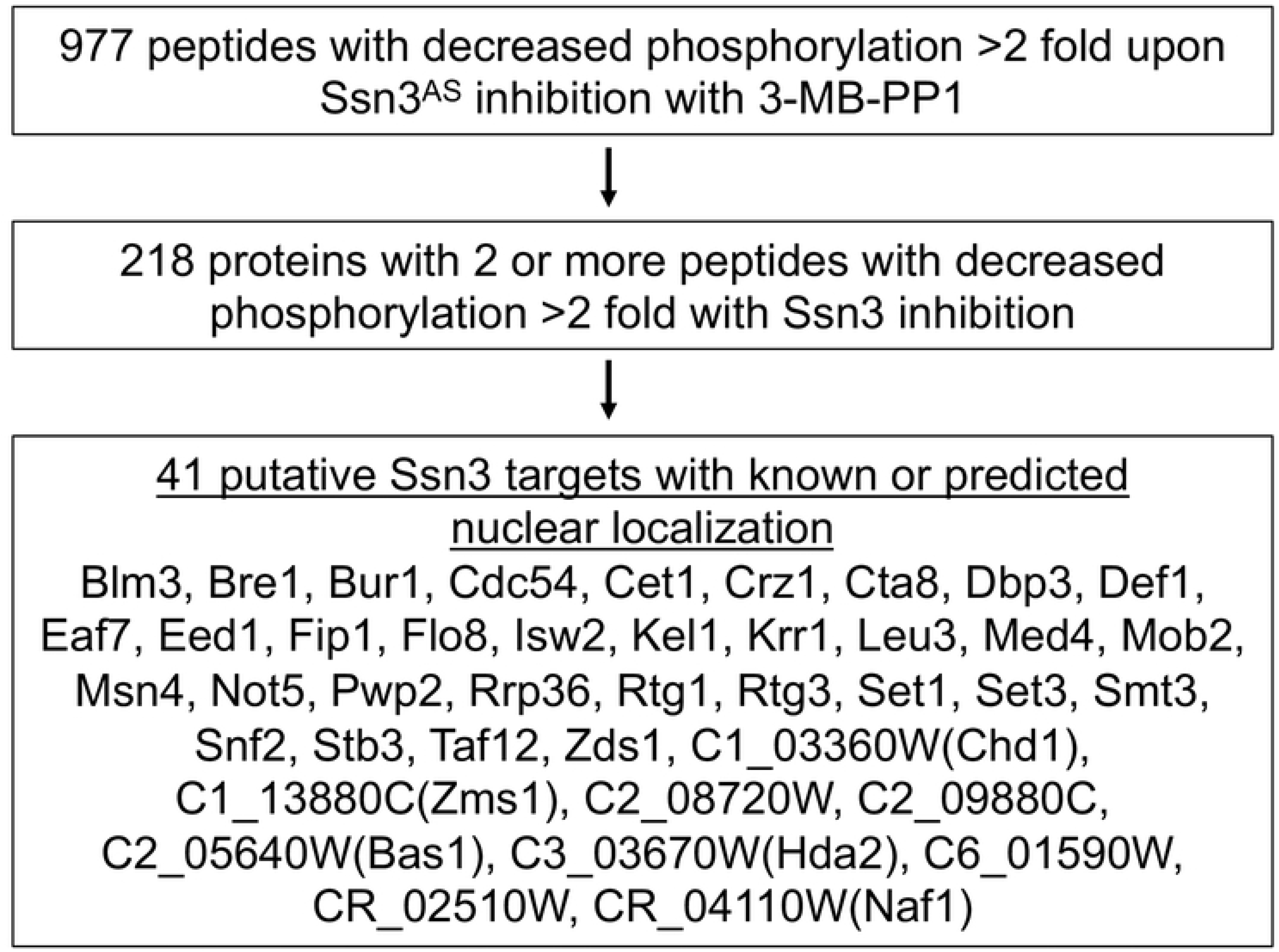
Overview of proteins for which two phosphopeptides were lower by >2 fold upon inhibition of Ssn3. Focusing on peptides that were significantly lower in Ssn3^AS^ treated with 3-MB-PP1 relative to control cultures (p<0.05 FDR-corrected), we found 977 peptides that were 2-fold lower upon 3-MB-PP1 inhibition of *ssn3^AS^* relative to changes in SC5314 WT. Two hundred and eighteen proteins had two more peptides that met these criteria, forty of which were predicted to have nuclear localization. Med4 is a validated *C. albicans* Ssn3 target. If an alias is available for an unnamed gene, it is shown in parentheses.

### Flo8 is required for Ssn3-dependent hyperfilamentation and hypha-specific gene expression

To determine whether there was a genetic interaction between *SSN3* and *FLO8* that accompanied the phosphoproteomic data above, we investigated the phenotype of single and double null mutants of these two genes. We previously reported that an *ssn3* null mutant forms wrinkled colonies in the presence of a metabolic inhibitor, pyocyanin, while the WT does not (12). While both the *ssn3* null strain and the WT formed smooth colonies on YNBA agar at 30°C (which is the same as YNBAG used above but without GlcNAc), only the ssn3Δ/Δ formed wrinkled colonies at 37°C. Similarly, the *ssn3*Δ/Δ strain, but not the WT, formed wrinkled colonies on solid YNBA medium with 110 mM added glucose (**Fig. 5A**). This wrinkled colony phenotype of the *ssn3*Δ/Δ mutant under the above conditions was abolished upon deletion of *FLO8* (*ssn3*Δ/Δ*flo8*Δ/Δ) (**Fig. 5A**). The *flo8*Δ/Δ mutant formed smooth colonies, like the WT, under all conditions. In liquid medium, we observed a similar epistatic relationship between *FLO8* and *SSN3*. Only the *ssn3*Δ/Δ strain, and not the WT, formed hyphae and the hyperfilamentation phenotype in *ssn3*Δ/Δ was dependent on *FLO8* (**Fig. 5B**). Expression levels of hypha-specific transcripts that were induced upon inhibition of Ssn3^AS^ (**Fig. 3**) were significantly higher in the *ssn3*Δ/Δ background compared to the WT, but not in the ssn3Δ/Δ*flo8*Δ/Δ background (**Fig. 5C**). In **Fig. 6**, we demonstrate the ability to complement the filamentation defects of the *flo8Δ*/Δ and *ssn3*Δ/Δ*flo8*Δ/Δ with the native *FLO8* allele, and this result is described in more detail below.

**Figure 5.**
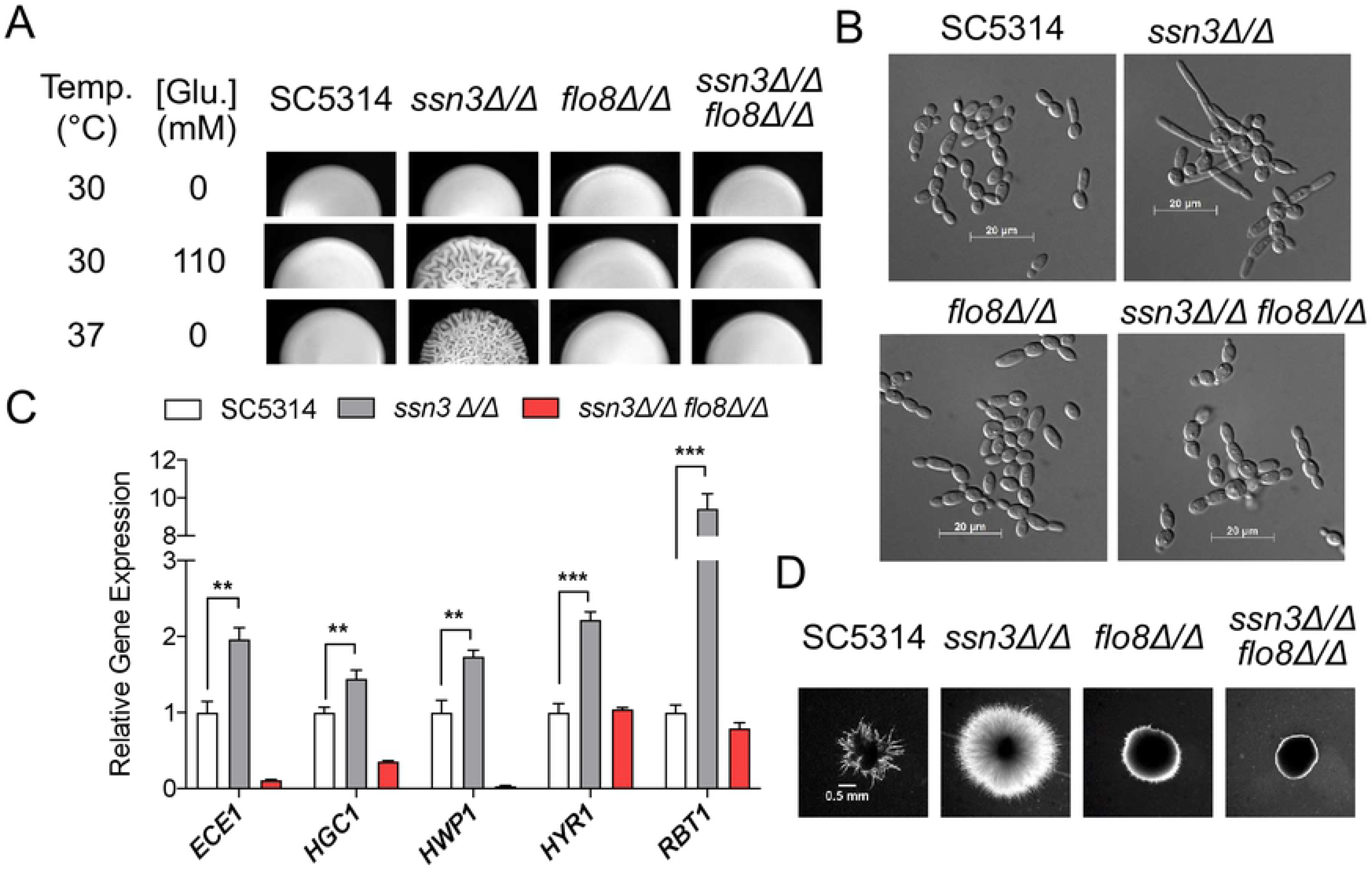
*SSN3* repression of filamentation is *FLO8* dependent. A. Colony morphology of a wild type *C. albicans* strain (SC5314), *ssn3Δ/Δ*, *flo8Δ/Δ* and *ssn3Δ/Δ flo8Δ/Δ* strains grown on YNBA agar medium alone or supplemented with 110 mM glucose at 30°C or on YNBA at 37°C. B. Cell morphology of strains tested in (A) after cells were grown in YNBN_2.5_AG_11_ at 30°C for 3 hours. C. NanoString analysis of indicated hypha-specific genes over-expression in the SC5314 wild type, and *ssn3*Δ/Δ, and *ssn3*Δ/Δ*flo8*Δ/Δ mutants. RNA was extracted from cells grown as indicated for (B) but for 75 minutes. Gene expression was represented by mean and standard deviation after normalization to Nanostring positive controls, and *TEF1* and *ACT1* reference transcripts; expression of each gene in the wild type strain (SC5314) was set to ‘1’; **, p<0.01 and ***,p<0.001. D. The embedded colonies of the same set of strains shown in (A) grown in YPS at 25°C.

**Figure 6.**
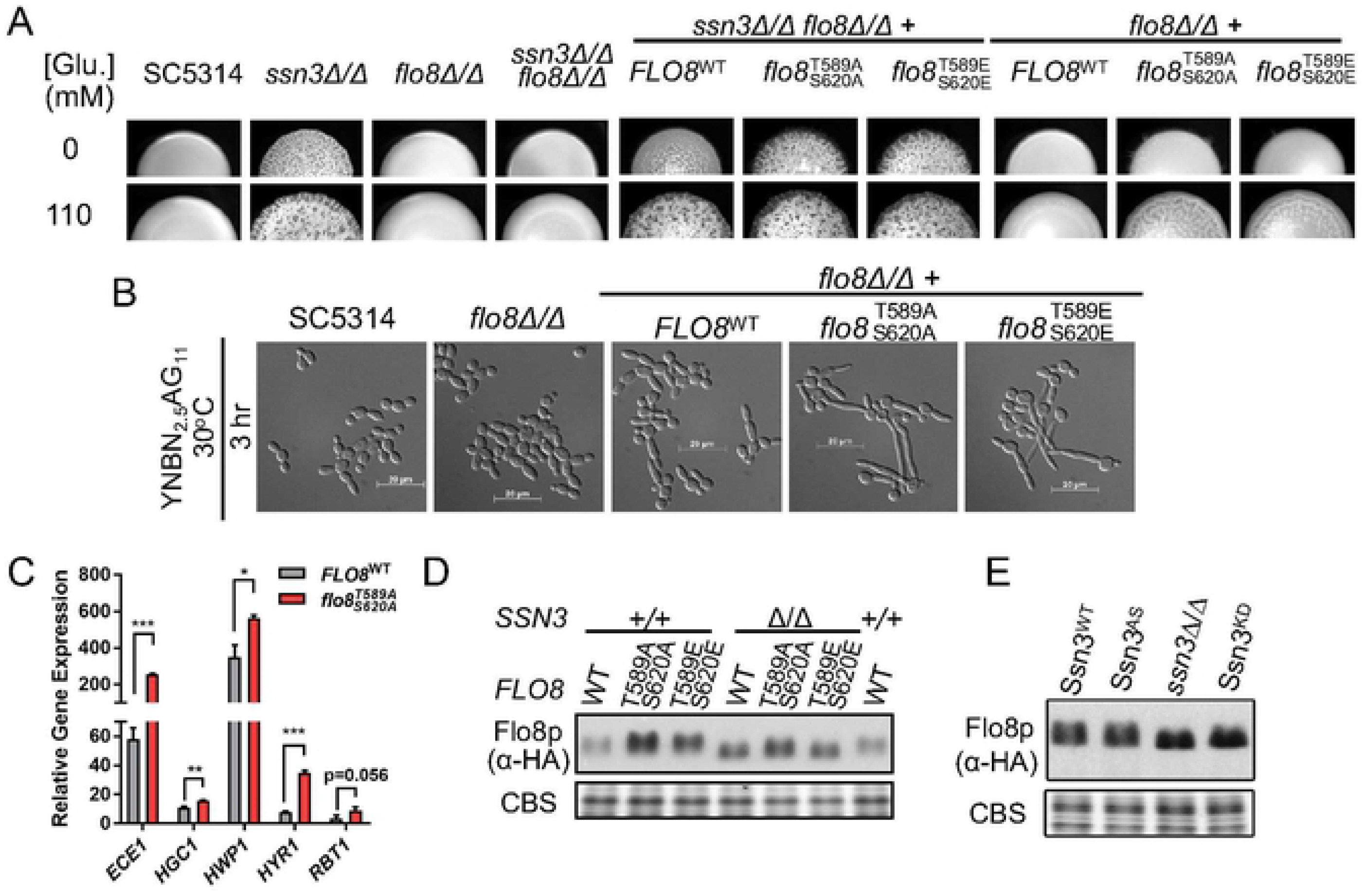
Residues identified as phosphorylation sites influence Flo8 function. A. SC5314 wild type, *ssn3Δ/Δ*, *flo8*Δ/Δ and *ssn3*Δ/Δ*flo8*Δ/Δ strains were imaged after growth as colonies on YNBA or YNBA+110 mM glucose at 37°C. *flo8*Δ/Δ and *ssn3*Δ/Δ*flo8*Δ/Δ expressing C-terminally 3XHA tagged Flo8^WT^, Flo8^T589A/S620A^ or Flo8^T589E/S620E^ were also included. B. Cell morphology of SC5314, *flo8*Δ/Δ, and *flo8*Δ/Δ expressing 3XHA-tagged Flo8^WT^, Flo8^T589A,S620A^ or Flo8^T589E,S620E^ after growth in YNBN_2.5_AG_11_ at 30°C for 3h. C. Gene expression in *flo8*Δ/Δ expressing 3XHA-tagged Flo8^WT^ or Flo8^T589A/S620A^ relative to *flo8*Δ/Δ after growth in YNBNAG for ~2h. Data show the mean and standard deviation from measurement on triplicate RNA samples. Gene expression in *flo8* mutant (not shown) was set to ‘1’. P-values (t-tests) were directly denoted or indicated by ‘*’ (p<0.05), ‘**’(p<0.01) or ‘***’ (p<0.001) to show statistically significant differences. D. Immunoblotting analysis with an α-HA antibody showing levels and gel mobility of native Flo8 with the C-terminal 3X-HA tag or variants with T589A,S620A or T589E,S620E substitutions; proteins were expressed in either *flo8*Δ/Δ or *ssn3*Δ/Δ *flo8*Δ/Δ backgrounds and were grown in YNBAG at 30°C to mid-log phase. E. Flo8-HA levels in strains with full Ssn3 activity, SC5314 (Ssn3^WT^) and *ssn3^AS^,* or without Ssn3 activity (*ssn3*Δ/Δ or *ssn3^KD^*).

We also found that the *ssn3*Δ/Δ strain was hyperfilamentous in comparison to the WT in embedded conditions, and that deletion of *FLO8* in the *ssn3*Δ/Δ background was able to suppress this phenomenon (**Fig. 5D**). Cao et al. (17) reported that *FLO8* was necessary for embedded filamentation and we reproduced this result (**Fig. 5D**). To determine whether *FLO8* is also required for embedded hyphal growth in *efg1Δ*/Δ, another strain that is hyperfilamentous under embedded conditions (28, 29), we generated a *flo8* and *efg1* null double mutant. We found that *FLO8* disruption suppressed embedded filamentation in the *efg1*Δ/Δ background (**Fig. S1A**).

### Loss of putative Ssn3-phosphosites in Flo8 leads to increased filamentation

To study the effects of Ssn3 activity on Flo8 protein, we complemented the *ssn3*Δ/Δ*flo8*Δ/Δ mutant with an allele that encodes a 3XHA-Flo8. This allele restored the hyperfilamentous *ssn3*Δ/Δ phenotype to the *ssn3*Δ/Δ*flo8*Δ strain (**Fig. 6A**). To further explore the effects of Flo8 phosphorylation by Ssn3, we generated an allele in which both T589 and S620, the sites identified in the phosphoproteomics analysis of the Ssn3AS bearing strain, were mutated to alanines (a phospho-incompetent residue) or glutamic acid (which also abolishes phosphorylation, but sometimes can serve as a phosphomimetic) to determine whether these residues play roles in the morphological phenotype of Cdk8 inhibition. Neither the WT or *flo8Δ/Δ* strain gave a wrinkled colony morphology on either medium (**Fig. 6A**) and, as expected, complementation of the *flo8*Δ/Δ with epitope tagged Flo8 did not change colony morphology. However, complementing the *flo8Δ/Δ* strain with either *flo8^T589A, S620A^* or *flo8^T589E, S620E^* led to a wrinkled colony phenotype in the presence 110 mM glucose, suggesting that the phosphorylation of Flo8, potentially via Ssn3, represses Flo8 activity and that the absence of these sites releases that repression (**Fig. 6A**). Consistent with the increased wrinkled colony phenotype in the *flo8*Δ/Δ strains with *flo8^T589A, S630A^* or *flo8^T589E, S620E^* relative to the strain with unmutated *FLO8*, both Flo8 variants also led to increased hyphal development in YNBAG (**Fig. 6B**). Because both *flo8^T589A, S620A^* and *flo8^T589E, S620E^* alleles conferred increased filamentation phenotypes consistent with high Flo8 activity in *flo8*Δ/Δ and *ssn3*Δ/Δ*flo8*Δ/Δ strains we concluded that the Flo8^T589E,S620E^ variant was not acting as a phosphomimetic, but rather both had phenotypes consistent with decreased negative regulation. To quantitatively assess the differences in activity of the morphology program attributed to the loss of putative Ssn3 phosphorylation sites, levels of hypha-associated transcripts were measured in the *flo8*Δ/Δ strains with either HA-tagged *FLO8* or *flo8^T589A, S620A^* (**Fig. 6C**). We found significantly increased expression levels of several core filamentation response genes in the strain with *flo8^T589A, S620A^* versus that with *FLO8*.

In multiple instances, phosphorylation by Ssn3 leads to decreased levels of transcription factors (6, 8, 15). Thus, we tested the hypothesis that Flo8-HA^T589A,S620A^ and Flo8-HA^T589E,S620E^ were present at higher levels than Flo8-HA. We found that indeed the two variants were present at higher relative levels compared to Flo8-HA in an *SSN3* wild-type background. Furthermore, Flo8-HA levels were higher in an *ssn3*Δ/Δ strain than in the *SSN3/SSN3* (+/+) strain which no significant differences in levels for the native and variant proteins. In order to address the slight difference in migration of Flo8-HA in the Δ/Δ strain backgrounds (**Fig. 6D**), we analyzed the migration of FLO8-HA in backgrounds with active Ssn3 (WT and *ssn3^AS^*) and strains lacking Ssn3 activity (*ssn3*Δ/Δ and *ssn3^KD^*) (**Fig. 6E**). We confirmed that in the absence of Ssn3 activity, Flo8 migration was slightly faster suggesting that other post-translational modifications (e.g. additional phosphorylations) are controlled by Ssn3. It is worth noting that one other Flo8 phosphosite was identified, but it did not reach the significance cutoff.

### Ssn3 metabolic hyperalkalinization phenotype depends on Flo8

Previous work has shown that the *C. albicans ssn3*Δ/Δ mutant differed from the WT in glycolysis and the utilization of amino acids (12). By analyzing the differentially abundant transcripts upon inhibition of Ssn3, we identified statistical enrichment of genes in KEGG pathways involved in amino acid metabolism and glycolysis (**Table S4**). These pathways could represent a potential connection between the *ssn3*Δ/Δ hyperalkalinization phenotype which is dependent on amino acid catabolism and concomitant release of ammonia (12). When grown on amino acid-containing YNBA, alkalinization of the agar medium, evident by the color change of the pH indicator, by the *ssn3*Δ/Δ mutant was less sensitive than WT to inhibition caused by increasing glucose concentrations which leads to production of acidic fermentation products. The previously observed alkalinization of the medium by the *ssn3Δ/Δ* strain (12) is consistent with increased alkalinization due to amino acid catabolism. To determine whether Ssn3 influences medium alkalinization through this mechanism, we constructed a *ssn3*Δ/Δ*stp2*Δ/Δ double mutant. Stp2 is a transcription factor involved in ammonia release from amino acid catabolism and is a downstream component of the SPS system (30). *STP2* disruption resulted in a defect in alkalinization at 5.5 mM glucose compared to WT and suppressed the hyperalkalinization phenotype of the *ssn3*Δ/Δ strain at both 27.5 mM and 110 mM glucose (**Fig. 7A**). This is consistent with a model in which the SPS amino acid sensing pathway and subsequent generation of ammonium through amino acid catabolism are primarily responsible for the hyperalkalinization phenotype of *SSN3* inactivation.

**Figure 7.**
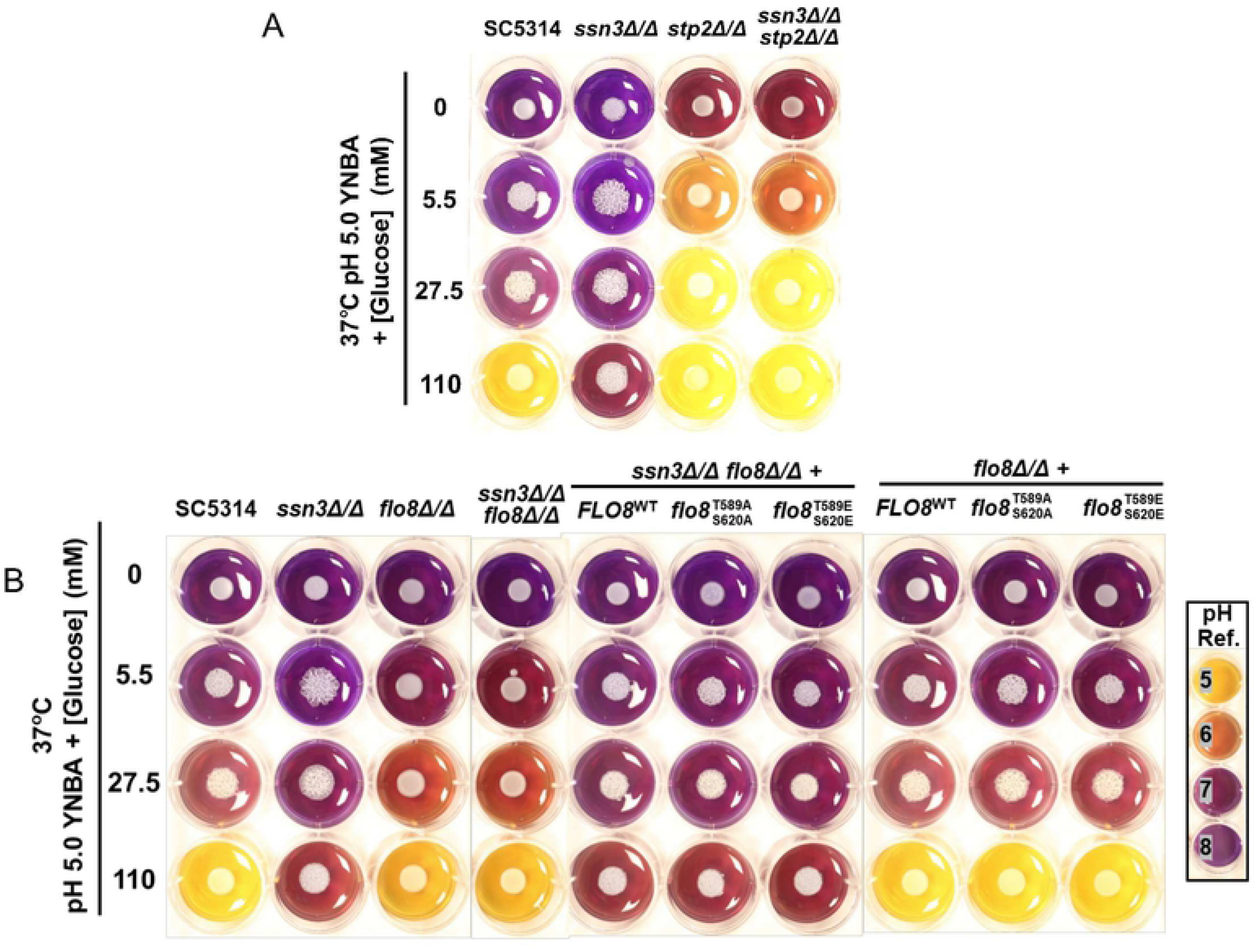
Medium alkalinization is affected by Ssn3 and Stp2. A. Comparison of medium pH by growth of SC5314, *ssn3*Δ/Δ, *stp2*Δ/Δ and *ssn3*Δ/Δ *stp2*Δ/Δ strains at 37°C. B. *ssn3* hyperalkalinization phenotype is dependent on *FLO8* at 37°C. SC5314, *ssn3*Δ/Δ, *flo8*Δ/Δ and *ssn3*Δ/Δ*flo8*Δ/Δ alone or expressing C-terminally 3XHA tagged Flo8^WT^, Flo8^T589A/S620A^ or Flo8^T589E/S620E^. Medium pH was assessed on YNBA with increasing concentrations of glucose and bromocresol purple as a pH indicator.

*FLO8* was required for the hyperalkalinization phenotype of the *ssn3*Δ/Δ strain and the phenotype was complemented by reintroducing *FLO8* (**Fig. 7B**), a result that paralleled the Flo8 requirement for hyperfilamentation in the *ssn3*Δ/Δ strain. In contrast, the *flo8*Δ/Δ strain did not differ from the WT in its effects on medium pH, suggesting that Flo8 effects on medium pH were associated with decreased Ssn3 activity (**Fig. 7A**). While complementation of the *ssn3*Δ/Δ*flo8*Δ/Δ double mutant with the wild-type *FLO8* allele restored the metabolic phenotypes, complementation with either *flo8^T589A,S620A^* or *flo8^T589E,S620E^* did not exacerbate the phenotype further. Similarly, complementation of the *flo8*Δ/Δ single mutant with the Flo8 variants did not generate changes in alkalinization compared to complementation with the native allele suggesting that Ssn3 effects on metabolism may require other factors directly or indirectly regulated by Ssn3.

## Discussion

In this work, we used strains with an analog-sensitive *ssn3* allele, a catalytically inactive *ssn3* allele and null mutants to assess how Ssn3 and its kinase activity regulate hyphal growth and metabolism. We identified 754 proteins with significant, differential phosphorylation (>2-fold, p<0.05) upon inhibition of Ssn3 during the yeast to hypha transition and they included Med4, a known Ssn3 substrate, and Flo8, a known regulator of hyphal growth (17, 18). We show that in a variety of conditions, the loss or decrease in Ssn3 activity leads to increased hypha formation and hypha-specific gene expression and that both phenotypes were dependent on Flo8. Through mutagenesis of the two Ssn3 phosphorylation sites (T589 and S620), we found that mutation of these sites was sufficient to increase filamentation and hypha-associated gene expression, supporting the model that Ssn3 is a negative regulator of Flo8. Interestingly, deletion of the *S. cerevisiae* histone methyltransferase *JHD2* and *SSN8* results in constitutive filamentous growth that requires the transcriptional positive regulator of invasive growth, *FLO8* (10). Our finding of 288 transcripts meeting our criteria for a statistically significant change in abundance upon the inhibition of Ssn3 is similar to the finding of Holstege and colleagues (7, 31) in which microarray analyses in *S. cerevisiae* found ~3% of genes changed by their criteria. The effects of the Ssn3^AS^ variant were modest compared to the effects of Ssn3 inhibition indicating the utility of this variant for the study of short term effects of Ssn3 inhibition.

In addition to Flo8, numerous of other transcriptional regulators of morphology have been described including Efg1, Cph1 (a homolog of *S. cerevisiae* Ste12, a known Ssn3 target), Tec1, Ndt80, and Ume6, and repressors like Tup1 and Nrg1 (32–39). Of these, Efg1 and Ndt80 were found to have significantly depleted phosphopeptides upon Ssn3 inhibition. We prioritized Flo8 because Wartenberg, *et al.* (14) recovered filamentation in a macrophage-evolved strain with *ssn3^R352N^* in an *efg1/cph1* null strain background, suggesting that regulators other than these were active upon changes in Cdk8 activity. Further, we showed that in embedded growth assays, the hyperfilamenation phenotype in both the *ssn3* and *efg1* null backgrounds required Flo8, underscoring the importance of Flo8 in Efg1-independent filamentation. Identification and functional characterization of phosphorylation sites and the understanding of their roles in Ssn3 regulation of activity merits further investigation. Flo8 has been implicated in the transcriptional regulation of true hyphal growth in *C. albicans*, with Flo8, and its binding partner Mss11, both having been described to directly bind the Hyphal Control Region in the promoter of *HWP1*, one of the most strongly differentially abundant transcripts in our transcriptomic data, and Flo8 is thought to be downstream of PKA (27, 40, 41). Ssn3 may play an important role in coordinating the response of multiple regulators during the induction and repression of filamentation. A recent study identified Ssn3 dependent phosphorylation of the transcription factor Ume6 and its degradation, under conditions of hypoxia and atmospheric CO_2_, as one way in which the kinase can impact filamentation (15). The absence of Ume6 phosphopeptides in our could be due to differences in kinetics of Flo8 and Ume6 or differences in the technologies used.

In addition to hyperfilamentation, wrinkled colony formation and medium alkalinization are phenotypes associated with *ssn3* deletion. As for hyperfilamentous growth, deletion of *FLO8* suppressed the hyperalkalinization phenotype of the *ssn3Δ/Δ* strain. Additionally, we were able to establish a role for Ssn3 upstream of the metabolic regulator Stp2 as deletion of *STP2* in the *ssn3* background was able to almost completely ablate the hyperalkalinization phenotype of the *ssn3Δ/Δ.* Stp2-mediated medium alkalinization is a mechanism by which *C. albicans* can regulate its morphology thus our findings are consistent with a model in which Ssn3 has both direct (Flo8-mediated) and indirect (Stp2-mediated) roles in determining morphology (42).

Several other pathways were identified as being influenced by Ssn3 in the proteomics studies including MAP kinase pathways, protein synthesis and chromatin regulation. It is interesting to note that studies of the human Cdk8 have also implicated it in regulating MAPK pathways (43). Notably, many of the phosphoproteins impacted by our inhibition of Ssn3 are not canonically nuclear, suggesting that Cdk8 may have a cytosolic role. This is in agreement with the work of Chen and Noble who identified a role of Cdk8 in the cytosolic phosphorylation of Sef1 (21). As an aside, we did not observe Sef1 phosphopeptides, a documented Ssn3 phosphotarget, and we speculate that this is due to the fact that our experiments were performed in an iron-replete medium which suppresses this phosphorylation event (21). We observed changes in proteins within MAP kinase cascades, their regulators, and effectors, and notable within that classification was the Hog1 MAP kinase pathway (**Table S3**). Among the depleted phosphopeptides in the *ssn3^AS^* strain were Ssk2 and Pbs2, the MAPKKK and MAPKK, respectively, of the Hog1 pathway, but Hog1 itself was not identified as being changed upon inhibition of Ssn3 activity. We also found differences in ribosomal biogenesis and protein synthesis. Lastly, we found that phosphopeptides involved in the remodeling of chromatin, such as the Set3 histone deacetylase, were depleted upon inhibition of Cdk8 kinase activity. This is similar to a phosphoproteomic study in human cells that identified elements of the NuA3 and NuA4 histone acetyltransferase complex amongst the depleted phosphopeptides upon inhibition of Cdk8 and the related kinase Cdk19 by cortistatin A (44). Set3 is also involved in morphological determination, with a *set3* null mutant displaying a hyperfilamentous phenotype (45). Additionally, it has been found that alterations in chromatin architecture participate in the interplay between Nrg1 and hypha-specific gene expression (39). The existence of a substantial suite of proteins involved in chromatin remodeling that show decreased phosphorylation with Ssn3 inhibition in the phosphoproteomic results is also a possible mechanism to account for the hyphal morphology phenotype of Ssn3 inhibition. A potential hypothesis is that Cdk8 phosphorylates hyphal transcriptional regulators like Flo8 acting in concert with phosphorylation of elements of the chromatin remodeling machinery, which promotes the formation of repressive chromatin structures at hypha-associated genes.

Unlike some other components of Mediator, which have pleiotropic effects on transcription, the role of the Cdk8 sub-module seems to be specific to certain developmental and nutrient regulated pathways across eukarya (22). This more specialized role has made Cdk8 a potential drug target for several diseases. For example, inhibitors of mammalian Cdk8 have been extensively explored as potential cancer therapies (46). As cancer therapeutics are often administered in the context of immunosuppression, it is important to understand the impact of these compounds on *C. albicans*, since inhibition of Ssn3 could potentiate or attenuate the virulence of this opportunistic pathogen.

## Materials and Methods

### Media and growth conditions

Strains were maintained on YPD plates, and overnight cultures for morphology and RNA seq were grown in YNB/1% glucose at 30°C in culture tubes on a roller drum. YPD plates were routinely streaked from glycerol stocks, and experiments were only conducted with overnight cultures from plates not more than 5 days old. YNB medium with 2% (w/v) casamino acids and 11 mM glucose (YNBAG) with or without 5 mM N-acetyl glucosamine (GlcNAc) was adjusted to pH 5.1 (morphology and RNA seq experiments) or pH 6.0 (phosphoproteomics) with concentrated hydrochloric acid, and filter sterilized. When indicated, the pH indicator bromocresol purple was included in the medium as described in (12). Additional medium, temperature, and incubation time details pertaining to morphology assessments can be found in the corresponding figure legends.

### Wrinkled colony and alkalinization assays

Overnight cultures (YPD) of each strain were washed once with water and diluted to OD_600_ 4. 8 μL of cell suspension was spotted onto the indicated medium. Images were typically taken after 2 days growth at 37°C or 3 days growth at 30°C. For alkalinization assays, 2X YNB based media was adjusted to pH 5.0 by HCl, filter sterilized and mixed with autoclaved 4% agar solution. Bromocresol purple (BCP) was added to 0.01% from a 0.1% aqueous stock into the media before 2 mL was aliquoted into each well of a 24-well plate. pH references were generated using YNB media buffered by phosphate buffer (20 mM) with known pH.

### Strain construction

All strains used in this study are listed in Table S1. Primers and plasmid sequences are available upon request. Strains expressing analog-sensitive and kinase-dead Ssn3 variants, referred to as *ssn3^AS^* and *ssn3^KD^*, respectively, were generated by transformation of SC5314 with a DNA fragment containing *ssn3-*F257G (*ssn3^AS^*) or *ssn3^D325A^* (*ssn3^KD^*) adjacent to the SAT-FLP cassette directed to the *SSN3* locus. These constructs were transformed alongside the *Candida*-optimized CRISPR/Cas9 machinery and a guide sequence targeting the nuclease to the *SSN3* open reading frame (47). Transformants were selected on YPD with 200 μg neourseothricin (GoldBio). Resistant colonies were patched onto YPD with 200 μg neourseothricin. No homozygous mutant transformants were identified, likely due to the guide sequence targeting the CRISPR/Cas9 nuclease to the mutant repair template. However, there were transformants likely for the intended point mutation (assayed with Sanger sequencing) which probably came about due to CRISPR-independent allelic replacement by homologous recombination in cells that did not also take up the Cas9 machinery. Amplification using primers that spanned the *SSN3* locus from DNA isolated from these heterozygotes revealed the *SAT1* marker was excised during outgrowth on YPD. Transformants were purified to single colonies, and neourseothricin sensitivity was confirmed by patching on YPD plates containing 200 μg per mL neourseothricin. Heterozygous point mutants underwent a second transformation with the same construct using the methods described above to generate homozygotes. Transformants in both rounds of transformation were confirmed by PCR amplifying a ≈970 base pair internal region of *SSN3* covering the region encoding residues 257 and 325 using primers Ssn3 Internal FWD and Ssn3 Internal Rev, and Sanger sequencing with the reverse primer. As in the first round, a number of these transformants had excised the resistance marker as observed by PCR during outgrowth on nourseothricin plates, and these sensitive homozygotes were purified to single colonies. Generation of subsequent knockout mutants was carried out using a previously described transient CRISPR-Cas9 system using a SAT-flipper selection marker (48). To generate double mutants, the *SAT1* cassette was recycled by inducing flippase expression in YP maltose (1% yeast extract, 2% peptone, 2% maltose) for 24 hours. Generation of HA-tagged flo8-bearing strains was similarly accomplished through a repair construct in which the desired allele was fused to the SAT-FLP marker.

### Mediator purification and in vitro kinase assays

Ssn8-tagged Mediator containing various Cdk8 alleles was purified and used for *in vitro* kinase assays with a GST-CTD substrate as previously described (20, 49). Amounts of the kinase were normalized by the signal on the FLAG tag.

### Analysis of C. albicans morphology

Morphological assessment was conducted in YNBNAG_11_ at 30°C. For the analysis of the effects of 3-MB-PP1 (the ATP analog used to inhibit analog-sensitive kinases), cells grown overnight at 30°C in YNB with 1% (w/v) glucose were pelleted by centrifugation and resuspended in 5 mL YNBNAG_11_ pH 5.1 containing 5 μM 3-MB-PP1 (EMD MILLIPORE) or DMSO as a vehicle control. The cells were then incubated at 30°C for 3h in culture tubes on a roller drum, fixed in formaldehyde, and morphology quantified from image captured using differential interference contrast microscopy. Cells were considered to be true hyphae if germ tubes had parallel sides and no invagination at the junction of the filament and the mother blastospore. A minimum of 175 cells per replicate were counted, and data presented represent three independent biological replicates conducted on three different days. Cultures were inoculated to an initial density of 1×10^7^ cells per mL.

### Analysis of the C. albicans transcriptome upon Ssn3 inhibition using RNA seq

Cells were grown as described above for morphological assessment, but the incubation time was reduced to decrease indirect effects of Ssn3 inhibition. Specifically, the time following drug exposure was reduced from three hours to one hour. The 5 mL cultures were collected one hour after 3-MB-PP1 addition, pelleted by centrifugation, snap frozen in liquid nitrogen, and RNA was isolated with the MasterPure™ Yeast RNA Purication Kit (Epicentre MPY03100). For RNA sequencing, 500 ng of total RNA was input into the Kapa mRNA HyperPrep kit (Kapa Biosystems, Wilmington, MA) and processed according to the manufacturer’s instructions. All 24 samples were multiplexed together into a single High Output 2×75bp run on a NextSeq 500 instrument (Illumina, San Diego, CA). Raw reads were mapped to the *C. albicans* genome SC5314 (version A21-s02-m09-r04, candidagenome.org), and normalized using EdgeR. KEGG enrichment analysis was carried out using KOBAS 2.0 (50).

RNA was extracted from frozen cell pellets as described (51). Data were presented after normalization by geometric mean of positive controls and geometric mean of *TEF1* and *ACT1* reads. Gene expression in a wild-type strain or a *flo8* mutant was set to ‘1’ as mentioned in the figure legends. The accession number for the data is GSE171859. During the review phase, RNA Seq data can be accessed at https://www.ncbi.nlm.nih.gov/geo/query/acc.cgi?acc=GSE171859 and the access token is mpsfuucitvktreh. The data will be publicly available upon acceptance for publication.

### Ssn3 phosphoproteomic experimental design

300 mL YNBNAG_11_ (pH 6.0) cultures of SC5314 and *ssn3^AS^* were inoculated at a density of OD_600_ of 0.01 and grown to OD_600_ of 4.5 in flasks with shaking at 30°C, then incubated for a subsequent 4.5 hours to ensure cells had entered stationary phase. Stationary phase cells were incubated in the presence of 5 μM 3-MB-PP1 for five minutes to enable drug entry into the cell. The cells were then concentrated and added to fresh, pre-warmed medium containing either drug or vehicle and incubated for 15 minutes at 30°C with shaking. Either DMSO vehicle or 5 μM 3-MB-PP1 was then added, after which cultures were incubated with shaking for five minutes at 30°C. Then, 1.2 L of prewarmed fresh YNBNAG_11_ (pH 6.0) medium was added as a 1.25X concentrate, and cells were incubated for an additional 15 minutes. The cells were then harvested by centrifugation and cell lysis by grinding under liquid nitrogen. All growth and incubation steps were conducted at 30°C, and the data represent the average of three independent replicates conducted on three separate days.

### Phosphoproteomic analysis

Yeast powder was lysed in ice-cold lysis buffer ((8 M urea, 25 mM Tris-HCl pH 8.6, 150 mM NaCl, phosphatase inhibitors (2.5 mM beta-glycerophosphate, 1 mM sodium fluoride, 1 mM sodium orthovanadate, 1 mM sodium molybdate) and protease inhibitors (1 mini-Complete EDTA-free tablet per 10 ml lysis buffer; Roche Life Sciences)) and sonicated three times for 15 sec each with intermittent cooling on ice. Lysates were centrifuged at 15,000 x *g* for 30 minutes at 4°C. Supernatants were transferred to a new tube and the protein concentration was determined using a BCA assay (Pierce-ThermoFisher Scientific). For reduction, DTT was added to the lysates to a final concentration of 5 mM and incubated for 30 min at 55°C. Afterwards, lysates were cooled to room temperate and alkylated with 15 mM iodoacetamide at room temperature for 45 min. The alkylation was then quenched by the addition of an additional 5 mM DTT. After 6-fold dilution with 25 mM Tris-HCl pH 8, the samples were digested overnight at 37°C with 1:100 (w/w) trypsin. The next day, the digest was stopped by the addition of 0.25% TFA (final v/v), centrifuged at 3500 x *g* for 30 minutes at room temperature to pellet precipitated lipids, and peptides were desalted on a 500 mg (sorbent weight) SPE C_18_ cartridge (Grace-Davidson). Peptides were lyophilized and stored at - 80°C until needed for future use.

### Phosphopeptide enrichment

Phosphopeptide purification was performed as previously described (52). Briefly, peptides were resuspended in 1.5 M lactic acid in 50% ACN (“binding solution”). Titanium dioxide microspheres were added and vortexed by affixing to the top of a vortex mixer on the highest speed setting at room temperature for 1 hour. Afterwards, microspheres were washed twice with binding solution and three times with 50% ACN / 0.1% TFA. Peptides were eluted twice with 50 mM KH_2_PO_4_ (adjusted to pH 10 with ammonium hydroxide). Peptide eluates were combined, quenched with 50% CAN / 5% formic acid, dried and desalted on a μHLB OASIS C_18_ desalting plate (Waters). Phosphopeptide enrichment was repeated once.

### TMT-labeling

Phosphopeptides were resuspended in 133 mM HEPES (Sigma) pH 8.5 and 20% acetonitrile (ACN) (Burdick & Jackson). Peptides were transferred to dried, individual TMT reagent (ThermoFisher Scientific), and vortexed to mix reagent and peptides. After 1 hr at room temperature, each reaction was quenched with 3 μl of 500 mM ammonium bicarbonate solution for 10 minutes, mixed, diluted 3-fold with 0.1% TFA in water, and desalted using C_18_ solid phase extraction cartridges (ThermoFisher Scientific). The desalted multiplexes were dried by vacuum centrifugation.

### Pentafluorophenyl-based Reversed Phase HPLC

Offline PFP-based reversed phase HPLC fractionation was performed as previously described (53). Briefly, phosphopeptides were fractionated using a Waters XSelect HSS PFP 2.5 μm 2.1 × 150 mm column on an Agilent 1100 liquid chromatography system, buffer A was 3% acetonitrile / 0.1% TFA, and buffer B was 95% acetonitrile / 0.1% TFA. Flow rate was 150 μl/min with a constant column temperature of 20°C. Phosphopeptides were fractioned using a 60-minute linear gradient from 8-45% acetonitrile and collected as 48 fractions between minutes 2 and 65. The 48 fractions were then combined into 24 total samples.

### TMT-based quantitative data analysis

TMT-labeled samples were analyzed on a Orbitrap Fusion (Senko, Remes et al. 2013) mass spectrometer (ThermoScientific) equipped with an Easy-nLC 1000 (ThermoScientific). Peptides were resuspended in 8% methanol / 1% formic acid across a column (45 cm length, 100 μm inner diameter, ReproSil, C_18_ AQ 1.8 μm 120 Å pore) pulled in-house across a 2 h gradient from 8% acetonitrile/0.0625% formic acid to 37% acetonitrile/0.0625% formic acid. The Orbitrap Fusion was operated in data-dependent, SPS-MS3 quantification mode (54, 55) wherein an Orbitrap MS1 scan was taken (scan range = 350 – 1500 m/z, R = 120K, AGC target = 2.5e5, max ion injection time = 100 ms), followed by ion trap MS2 scans on the most abundant precursors for 4 seconds (max speed mode, quadrupole isolation = 0.6 m/z, AGC target = 4e3, scan rate = rapid, max ion injection time = 60 ms, minimum MS1 scan signal = 5e5 normalized units, charge states = 2, 3 and 4 included, CID collision energy = 33%) and Orbitrap MS3 scans for quantification (R = 15K, AGC target = 2e4, max ion injection time = 125 ms, HCD collision energy = 48%, scan range = 120 – 140 m/z, synchronous precursors selected = 10). The raw data files were searched using COMET with a static mass of 229.162932 on peptide N-termini and lysines and 57.02146 Da on cysteines, and a variable mass of 15.99491 Da on methionines and 79.96633 Da on serines, threonines and tyrosine against the target-decoy version of the respective FASTA database (UniProt; www.uniprot.org) and filtered to a <1% FDR at the peptide level. Quantification of LC-MS/MS spectra was performed using software developed in house. Phosphopeptide intensities were adjusted based on total TMT reporter ion intensity in each channel and log_2_ transformed. P-values were calculated using a two tailed Student’s t-test assuming unequal variance.

## Acknowledgments

Research reported in this publication was supported by National Institute of Health (NIH) through the National Institute of Allergy and Infectious Disease grants AI127548 to D.A.H., AI115253 to L.C. M., and T32 AI007519 to J.M.H. and National Institute for General Medical Science grant R35 GM119455 for A.N.K.. This work was also supported by P30-DK117469 for the Applied Bioinformatics and Biostatistics Core. Sequencing services and specialized equipment was provided by the Genomics and Molecular Biology Shared Resource Core at Dartmouth supported by NCI Cancer Center Support Grant 5P30CA023108-37. Equipment used was supported by the NIH IDeA award to Dartmouth BioMT P20-GM113132. The content is solely the responsibility of the authors and does not necessarily represent the official views of the NIH.

---

**Figure S1.** *FLO8* is required for hyperfilamentation in an *efg1*Δ/Δ mutant in embedded conditions and in colony biofilms. A. SC5314 wild type parental strain, *flo8*Δ/Δ, *flo8*Δ/Δ complemented with *FLO8*, *efg1*Δ/Δ, *efg1*Δ/Δ*flo8*Δ/Δ, *efg1*Δ/Δ*flo8*Δ/Δ+*FLO8* as embedded colonies in YPS agar 23°C or YNBA agar with 110 mM glucose at 30°C. The *efg1Δ*/Δ is hyperfilamentous in embedded colony conditions and the filamentation is dependent on *FLO8*.

## Supplementary Data

**Table S1.** Strains used in this work.

**Table S2.** RNA seq analyses to determine the effects of specific inhibition of ssn3-as by 3-MB-PP1 (drug). Sheets one and two compare the effect of drug versus vehicle in SC5314 and ssn3as, respectively.

**Table S3.** Phosphoproteomic analysis of the effect of 3-MB-PP1 on ssn3^as^ versus SC5314. Sheet one provides an annotated summary of phosphopeptides statistically significantly altered by drug treatment of compared to drug treatment of SC5314. Peptides highlighted in red indicate phosphorylation of a serine or threonine with a proline in the +1 or +2 position. Sheet two provides the data with each replicate of the triplicate experiment parsed individually.

**Table S4.** KEGG pathway enrichment analysis of transcripts altered in abundance in the *ssn3^as^* strain by 3-MB-PP1 treatment vs vehicle.

